# Machine learning reveals the diversity of human 3D chromatin contact patterns

**DOI:** 10.1101/2023.12.22.573104

**Authors:** Erin N. Gilbertson, Colin M. Brand, Evonne McArthur, David C. Rinker, Shuzhen Kuang, Katherine S. Pollard, John A. Capra

## Abstract

Understanding variation in chromatin contact patterns across human populations is critical for interpreting non-coding variants and their ultimate effects on gene expression and phenotypes. However, experimental determination of chromatin contacts at a population-scale is prohibitively expensive. To overcome this challenge, we develop and validate a machine learning method to quantify the diversity 3D chromatin contacts at 2 kilobase resolution from genome sequence alone. We then apply this approach to thousands of diverse modern humans and the inferred human-archaic hominin ancestral genome. While patterns of 3D contact divergence genome-wide are qualitatively similar to patterns of sequence divergence, we find that 3D divergence in local 1-megabase genomic windows does not follow sequence divergence. In particular, we identify 392 windows with significantly greater 3D divergence than expected from sequence. Moreover, 26% of genomic windows have rare 3D contact variation observed in a small number of individuals. Using *in silico* mutagenesis we find that most sequence changes to do not result in changes to 3D chromatin contacts. However in windows with substantial 3D divergence, just one or a few variants can lead to divergent 3D chromatin contacts without the individuals carrying those variants having high sequence divergence. In summary, inferring 3D chromatin contact maps across human populations reveals diverse contact patterns. We anticipate that these genetically diverse maps of 3D chromatin contact will provide a reference for future work on the function and evolution of 3D chromatin contact variation across human populations.

## 1 Introduction

Genetic and transcriptomic variation within and between human populations is extensive (1000 Genomes Project Consortium et al., 2015; Alemu et al., 2014; Ho et al., 2008; Mallick et al., 2016; Novembre et al., 2008; Storey et al., 2007). Understanding the implications of non-coding genetic variation and the causes of transcriptional variation remains challenging, particularly regarding the role of non-coding variation in generating phenotypic diversity across human populations. Therefore, comprehending the diversity and impacts of non-coding variation is crucial for advancing understanding of gene expression regulation and phenotypic variance.

The three-dimensional spatial organization of chromosomes within the nucleus, known as 3D chromatin contact, influences gene expression regulation through enhancer modulated transcription (Tang et al., 2015; Tolhuis et al., 2002). Experimental data has provided valuable insights into chromatin structure and interactions within the nucleus, such as data from the 4D Nucleome Project (Dekker et al., 2017, 2023). For example, disruption of the structural organization and contacts of distal regulatory elements within the genome has been linked to complex diseases and genomic rearrangements, such as those observed in certain cancers (Maurano et al., 2012; Roix et al., 2003; Zhang et al., 2012). Nonetheless, our knowledge of the breadth of 3D genome variation across genetically diverse human populations is still limited.

Previous studies have shown that 3D chromatin contact varies both within and among populations (Li et al., 2023; McArthur et al., 2022). Experimental determination of chromatin interactions at a population scale is expensive, especially at high enough spatial resolution to reveal differences in contacts between specific regulatory elements. This has limited the extent to which chromatin contact diversity has been studied across human populations. However, recent advances in machine learning methods have allowed for the prediction of 3D genome chromatin contact maps from DNA sequences (Fudenberg et al., 2020; Schwessinger et al., 2020; Zhou, 2021). These methods predict 3D chromatin contact based solely on sequence information, offering a promising approach to computationally study 3D genome diversity.

In this study, we used Akita (Fudenberg et al., 2020), which is a flexible convolutional neural network that requires only DNA sequence information as input, to predict 3D contact maps for 2,457 diverse human individuals. We compared these contact maps between individuals and to the predicted map of an inferred ancestral hominin genome sequence. This revealed regions with significant differentiation in contact maps between populations that may contribute to phenotypic differences. We found that 3D contact divergence genome-wide follows similar patterns as sequence divergence and that pressure to maintain 3D contact patterns has broadly constrained sequence evolution. However, 3D contact diversity is very different from sequence diversity at the local (1 Mb) scale. We also identified loci with significant variation in 3D chromatin contacts that associate with high evolutionary conservation and binding sites of CTCF—a transcription factor and critical determinant of 3D genome structure. Our results establish the baseline distribution of 3D chromatin contact and variation in diverse populations. They also provide context in which to interpret new human 3D chromatin contact data and the effects of variants identified in disease cohorts on 3D chromatin contact.

## 2 Results

### 2.1 Accurate prediction of 3D contact maps for diverse individuals

To quantify variation in the 3D genome of modern humans, we predicted chromatin contact maps for 2,457 unrelated individuals from the 1000 Genomes Project (1KGP) data (1000 Genomes Project Consortium et al., 2015) using Akita (**Figure 1A**) (Fudenberg et al., 2020). Akita takes an approximately 1 Mb window of DNA sequence as input and outputs local 3D contact patterns for the input region at 2,048 bp resolution. We generated pseudo-haploid genome sequences for each individual by inserting all their single nucleotide variants (SNVs) into the hg38 human reference sequence. We divided the genome into 1 Mb sliding windows, overlapping by half, and retained windows with 100% sequence coverage from the hg38 reference genome (N=4,873). We then used Akita to predict local chromatin contacts genome-wide for diverse individuals from five continental populations encompassing 26 unique sub-populations distributed across the globe (1000 Genomes Project Consortium et al., 2015).

**Figure 1:**
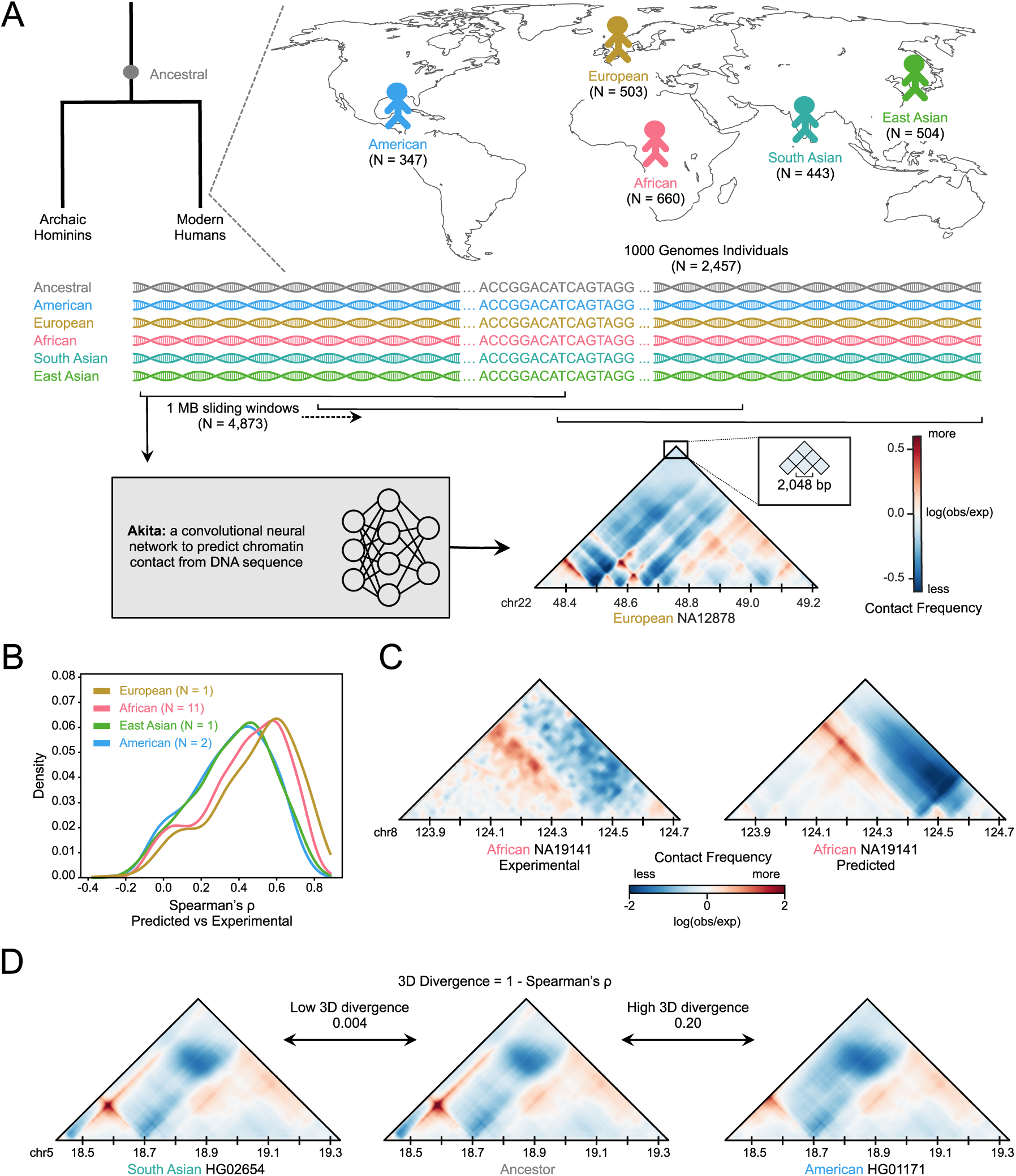
Strategy for investigating 3D chromatin contact patterns in diverse human populations. **(A)** Schematic of the generation of genome-wide 3D contact maps for 2,457 unrelated individuals from the 1000 Genomes Project. Akita is a deep neural network that takes approximately 1 Mb of DNA sequence as input and generates a 3D contact map of the window. The map consists of contacts for all pairs of 2,048 bp regions within the window. We applied Akita to the DNA sequence of each individual in sliding windows overlapping by half across the genome. We discarded windows without full sequence coverage in the hg38 reference sequence, resulting in 4,873 windows. We also applied Akita to an inferred human–archaic hominin ancestral sequence (Wohns et al., 2022). **(B)** Density of Spearman correlations between experimental and predicted contact maps at 10 kb resolution for windows in the Akita held-out test set of 413 windows across 15 individuals from 4 populations. This includes a European individual (GM12878) used as part of the Akita training data as a benchmark. The strong performance on African individuals suggests that Akita is accurate across populations. The lower performance on the East Asian and admixed American individuals is likely due to lower resolution of their experimental maps (**Figure S1**). **(C)** Example experimental and predicted maps for a representative window on chromosome 8 (chr8:123,740,160-124,788,736) from an African individual. **(D)** Example predictions and comparisons of 3D chromatin contact maps between pairs of individuals on chromosome 15 (chr15:18,350,080-19,398,656). To quantify “3D divergence”, we calculated the Spearman correlation coefficient over the corresponding cells for a given pair of maps subtracted from 1. Low 3D divergence scores indicate high similarity between contact maps and high divergence scores indicate low similarity between maps.

In order to confirm that Akita performs well on diverse individuals, we compared experimentally determined to predicted maps from 11 African, 2 American, 1 East Asian, and 1 European individual with Hi-C from the 4D Nucleome Project (4DN) (Dekker et al., 2017). We focused on held-out windows from the Akita test set and scaled predictions to 10 kb resolution to be roughly comparable to the lower resolution of the experimental contact maps. The European individual (NA12878) was the basis for the GM12878 lymphoblastoid cell line which was used in the training of Akita. Its Hi-C library was also sequenced to the highest coverage **Figure S1**. Thus, it serves as an upper bound on the expected performance. Our predictions for the 11 African individuals (mean Spearman’s *ρ* = 0.43) were only slightly less accurate than what was observed for Europeans (*ρ* = 0.48) (**Figure 1C**). While the East Asian (*ρ* = 0.37) and American (*ρ* = 0.36) accuracy ranges are somewhat lower, we believe this is due to low resolution and sequencing depth of the available experimental maps for these individuals. To test this, we correlated filtered read count (retrieved from 4DN Data Portal) with Akita prediction accuracy and found a correlation (*R*^2^ = 0.25, **Figure S1**). Visual checks verify that the predicted and experimental contact maps share key patterns (**Figure 1D**). These results confirm that Akita has learned to predict 3D contact maps in a way that is not specific to any single human or population and thus can be applied across diverse samples.

To compare predicted contact maps for the same genomic region from two individuals, we define the 3D divergence of a window to be one minus the Spearman correlation (1 – *ρ*) between the two maps (**Figure 1B**). We use two comparison schemes throughout the work. The first compares contact maps for each modern individual to contact maps predicted from the inferred sequence of the common ancestor of modern humans and archaic hominins (Wohns et al., 2022). The second compares contact maps between pairs of modern individuals using a representative subset of the cohort for computational efficiency (N = 130).

### 2.2 3D genome divergence differs from sequence divergence

To explore how changes in 3D chromatin contacts relate to DNA sequence changes, we compared sequence divergence from the ancestral sequence with 3D divergence from the ancestral map for each individual. Genome-wide average sequence and 3D divergence were only moderately correlated (**Figure 2A**; R^2^ = 0.31). variation in correlation strength by population suggests that the relationship between sequence divergence and 3D genome organization is complex and may be influenced by population-specific factors. The strength of this correlation shows that, while they are related, 3D divergence provides different information than overall sequence divergence. These results additionally suggest that 3D chromatin contact variation is much more constrained than sequence variation.

**Figure 2:**
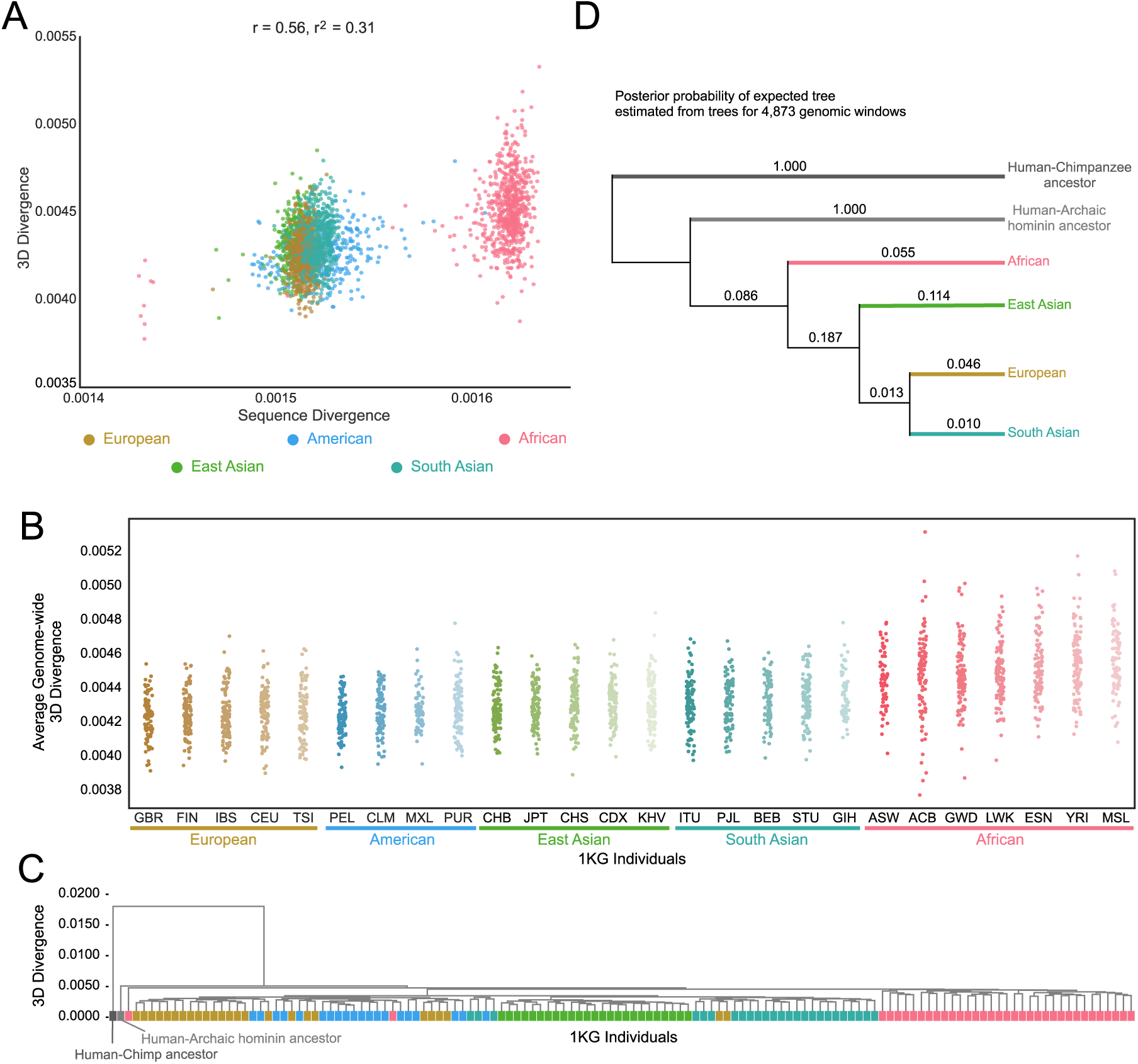
Genome-wide 3D divergence follows known population structure. **(A)** The relationship between genome-wide average sequence and 3D divergence from the human–archaic hominin common ancestor for each 1KG individual. **(B)** Genome-wide average 3D divergence for each 1KG individual, stratified by continental and sub-continental populations. Color indicates super-population and hue indicates subpopulation. **(C)** Genome-wide clustering of 1KG individuals with the inferred human-archaic hominin and human-chimpanzee ancestors using average genome-wide 3D divergence. Color indicates super-population. **(D)** Branch support (posterior probability) for the population tree inferred from 1KG sequences estimated using ASTRAL (Zhang et al., 2018) from the topologies of trees constructed for each window based on 3D divergence. Color indicates super-population.

### 2.3 African populations have the highest predicted 3D genome diversity

We quantified levels of 3D divergence genome-wide between diverse modern humans and the inferred hominin ancestor. We calculated 3D divergence from ancestral sequence for each window and took the mean across all genomic windows for each individual. Based on the higher sequence diversity of African populations (1000 Genomes Project Consortium et al., 2015), we expected that African populations would also have higher predicted 3D divergence from the ancestral state than in other populations.

Mean genome-wide 3D divergence varies significantly among populations (**Figure 2B**; Kruskal-Wallis: *P* = 2.34 *×* 10*^−^*^145^). African individuals have significantly greater mean divergence (0.0045) than individuals from all other populations (post-hoc Conover: *P <* 1.35 *×* 10*^−^*^57^), and non-African populations have on average 5% lower 3D divergence. While this is consistent with patterns of sequence divergence, the size of the difference is smaller; non-African individuals have approximately 20% fewer SNVs on average (1000 Genomes Project Consortium et al., 2015). These results suggest that while related, much of sequence variation has little impact on 3D chromatin contact.

### 2.4 Most variation in 3D chromatin contact patterns is shared across populations

To explore the similarity of 3D contact maps within and between humans from diverse populations, we hierarchically clustered five representative 1KG individuals from each of the 26 sub-populations (N = 130) based on their pairwise 3D divergence. Averaging over all 4,873 genomic windows, individuals grouped roughly by population of origin (**Figure 2C**). In contrast, clustering each window of the genome separately for these individuals revealed a diversity of relationships that did not follow global population relationships expected from sequence divergence patterns. To summarize the patterns across windows, we computed the posterior probability of the tree derived from sequence relationships based on all of the window-specific 3D divergence trees using ASTRAL (Rabiee et al., 2019; Zhang et al., 2018). Branches leading to modern populations are not strongly supported, reflecting the sharing of contact patterns between populations (**Figure 2D**). While the population branches are not well supported, the branches leading to inferred human-archaic hominin and human-chimpanzee ancestors each have posterior probabilities of 1.00. These results collectively indicate that 3D genome variation among modern humans is typically not stratified by population in any given genomic locus, but population structure emerges over longer evolutionary time periods and genomic distances.

### 2.5 3D genome divergence is highest in regions with the lowest functional constraint

The previous sections largely focused on patterns of 3D divergence summarized at the genomewide level. To quantify local patterns of 3D divergence along the genome for each genomic window, we computed the 3D divergence of each 1KG individual from the ancestral map. The mean 3D divergence in each window is highly variable across the genome, with many distinct peaks and valleys in both the mean (**Figure 3A**) and standard deviation (**Figure S2**). The majority of the top 10% most divergent windows are shared by all five continental populations (**Figure S3**). Taken together, these results demonstrate that some windows harbor greater 3D genome divergence, while others exhibit only slight variations on a widely shared contact pattern.

**Figure 3:**
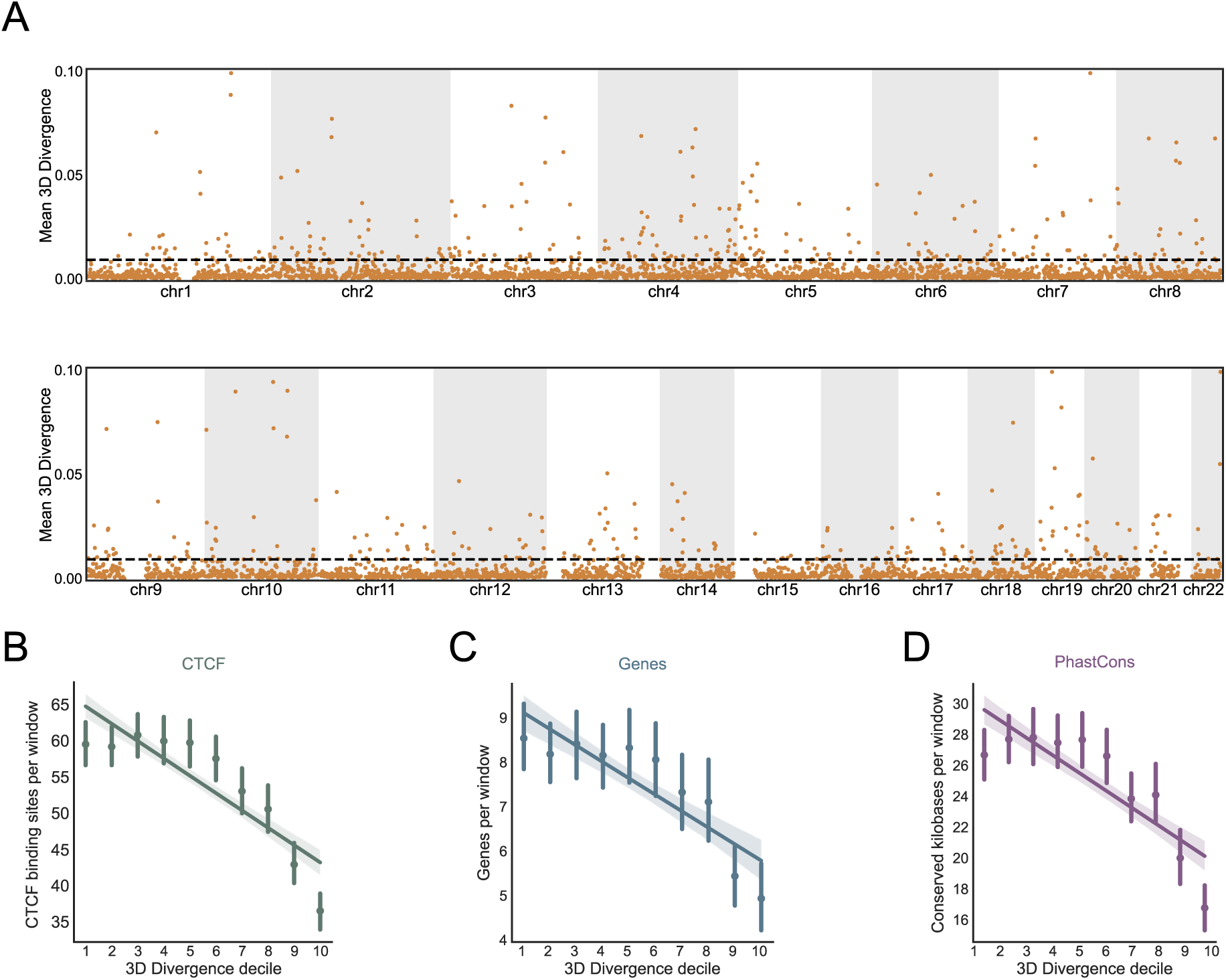
3D divergence is variable across the genome and highest in less functional regions. **(A)** Mean 3D divergence from the human-archaic hominin ancestor across 2,457 individuals from 1KG for each of the 4,873 genomic windows. Each point represents the mean divergence of all individuals from the ancestral genome for a single genomic window. The dotted line indicates the top 10% of 3D divergence. Divergences greater than 0.10 are plotted at 0.10 to aid visualization. **(B)** Average number of CTCF binding sites per window stratified by decile of mean 3D divergence from the hominin ancestor (bin 1 has the lowest divergence and 10 highest). Bars indicate bootstrapped 95% confidence intervals. CTCF peaks come from merging CTCF ChIP-seq peaks across all cell types from the ENCODE Consortium. **(C)** Average number of genes per window from GENCODE version 24 in each 3D divergence decile. Visualized as in **B. (D)** Average PhastCons 100-way conserved bases (in kb) per window in each 3D divergence decile. Visualized as in **B.**

Given the variation in 3D divergence from ancestral across the genome, we tested whether the level of 3D divergence associates with functional annotations or evolutionary sequence conservation between species. We stratified the genomic windows into 10 equal-size bins based on increasing 3D divergence and quantified the gene count, CTCF site count (ENCODE Project Consortium et al., 2020), and PhastCons 100-way conserved elements (Siepel et al., 2005) distributions for each decile.

Increasing 3D divergence consistently correlates with decreases in sequence identity, gene content, CTCF binding sites and PhastCons conserved bases (**Figures 3B–3D**). We also considered SPIN state (Kamat et al., 2023; Wang et al., 2021) predictions and repeat element annotations (Smit, 1999; Smit et al., 1996–2010), but did not observe an overall trend in SPIN state or repeat element prevalence (**Figures S4A**, **S4B**). However, “Lamina” and “Lamina-like” SPIN states are more prevalent in higher 3D divergence windows, while active states are less prevalent (**Figure S4A**). These results suggest regions with many functional elements or high sequence conservation have less 3D divergence, while those with less functional activity and conservation are more tolerant of variation in 3D contacts, meaning that 3D chromatin contacts may contribute to evolutionary pressures on sequence divergence.

### 2.6 3D chromatin contact constrains sequence evolution

Next, we explored whether the amount of 3D divergence between humans and the human-archaic hominin ancestor is more or less than expected given the observed sequence divergence. To estimate the expected 3D divergence distribution for each window, we generated 500 sequences with the number of sequence variants from the ancestral matched to the distribution across 1KG individuals and applied Akita to predict the resulting 3D genome divergence (McArthur et al., 2022). We preserved the tri-nucleotide context of all variants in each window for each sequence to account for variation in the mutation rate across sequence contexts. For each window, we compared the observed 3D divergence with the expected 3D divergence from the 500 sequences with the matched level of nucleotide divergence. If the pressure to maintain 3D chromatin contact patterns does not influence sequence divergence, the observed 3D divergence would be similar to the expected 3D divergence. If the observed 3D divergence deviates from the expected based on sequence divergence, more divergence would suggest positive selection on variants causing 3D differences, while less divergence would suggest negative selection on variants causing 3D differences.

The observed 3D divergence is significantly less than expected based on sequence divergence (**Figure 4**). 88.7% of windows have less 3D divergence than expected based on their observed sequence differences (binomial test *P <* 2.23 *×* 10*^−^*^308^). Genome-wide, the mean expected 3D divergence is 70% higher than the observed 3D divergence (*t*-test *P* = 1.68*×*10*^−^*^74^). This suggests that pressure to maintain 3D genome organization constrained sequence divergence in recent human evolution. This aligns with previous studies that demonstrated depletion of variation at 3D genome-defining elements (e.g., TAD boundaries, CTCF sites) (Fudenberg and Pollard, 2019; McArthur and Capra, 2021), and it specifically implicates maintenance of 3D chromatin contacts as a driver of sequence constraint.

**Figure 4:**
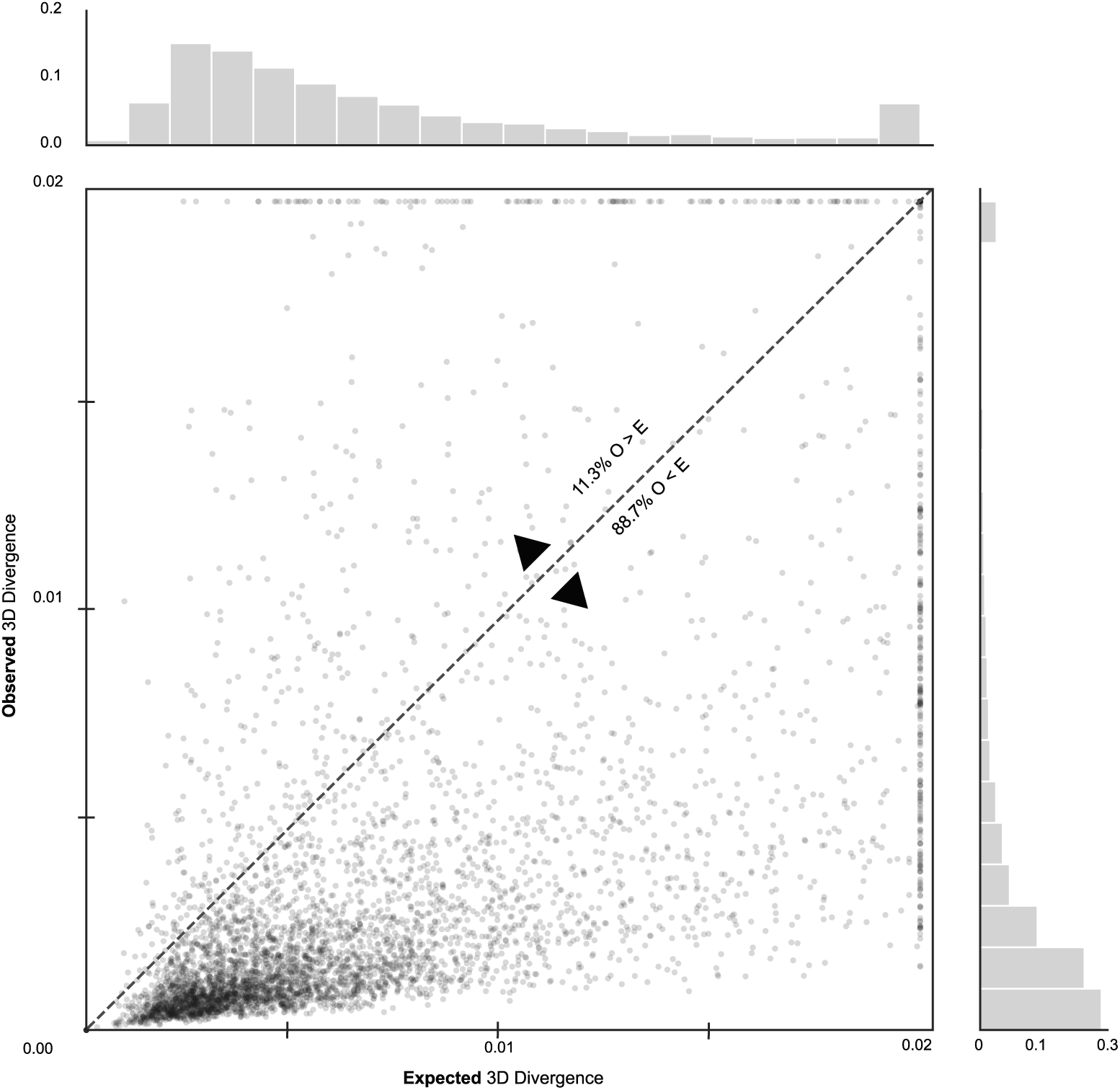
3D divergence is lower than expected in 89% of genomic windows, but 392 have significantly greater divergence than expected. Mean observed 3D divergence between 1KG individuals and the human-archaic hominin ancestor compared to 3D divergence expected based on amount of sequence variation. The expected 3D divergence distribution for a window is based predicted 3D genome organization for 500 simulated sequences for each window (Methods). Points above the line are windows more divergent than expected, which suggests more observed variants that alter 3D divergence than expected. Points below are windows less divergent than expected, which suggests constraint on sequence variation to maintain 3D chromatin contact. Observed 3D divergence is significantly less than the mean expected 3D divergence based on sequence (O < E: 88.7% of N = 4322 windows below the diagonal, binomial-test *P <* 2.23 *×* 10*^−^*^308^). The mean expected 3D divergence is on average 70-times higher than the observed 3D divergence (t-test *P* = 1.68 *×* 10*^−^*^74^). In contrast, 392 windows have distributions of observed 3D divergence significantly greater (t-test *P <*= 0.05) than the 3D divergence expected based on sequence divergence (O > E). 3D divergence scores greater than 0.02 are plotted at 0.02 for visualization.

### 2.7 392 windows have significantly greater 3D divergence than expected

Even though most windows have lower 3D divergence than expected, we found 392 windows in which the distribution of observed 3D divergence is significantly greater (*t*-test *P <*= 0.05) than the 3D divergence expected based on sequence divergence (**Figure 4**). These windows usually have many individuals with high 3D divergence, and we refer to them as “3D divergent windows”. For example, a 3D divergent window on chromosome 1 (chr1:88,604,672-89,653,248) exhibits a multi-modal 3D divergence distribution: a portion of the individuals fall within the expected divergence levels and a portion are much more divergent (**Figure 5A**). When stratified by populations, the vast majority of 3D divergent windows are divergent in all five continental populations, followed by African-specific divergent windows and divergent windows specific to non-African populations (**Figures S5**, **5B**). In the example window, our predictions show a group of individuals with a notable loss in contact compared to the other group of individuals with contact maps similar to the ancestral map (**Figures 5C**, **5D**). Using experimental data from the 4DN we confirmed the presence of both predicted patterns in experimental Hi-C maps (**Figures 5C**, **5D**). These results demonstrate that some genomic windows have substantial 3D genome variation within human populations.

**Figure 5:**
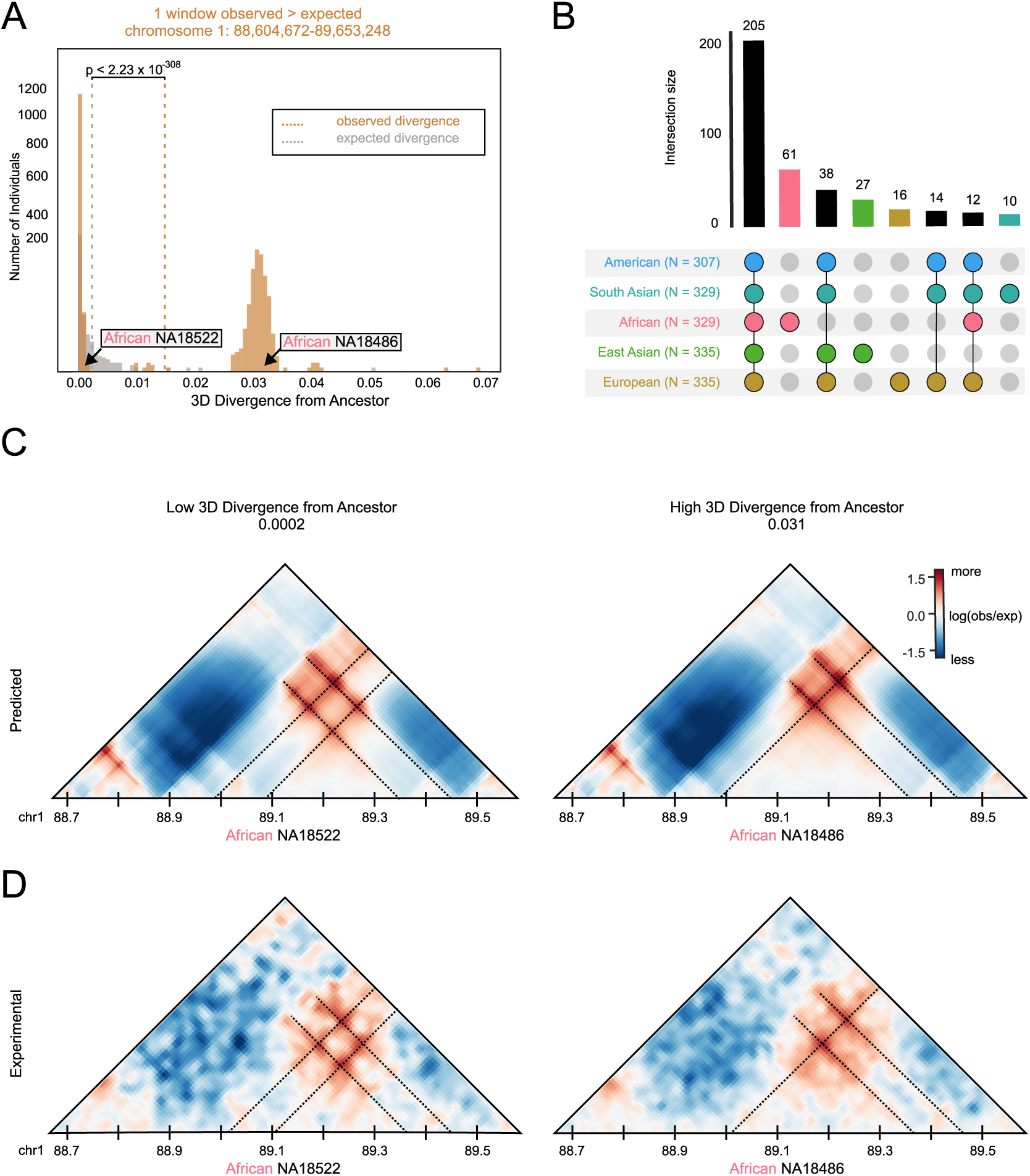
Experimental Hi-C data confirms predicted contacts in highly divergent windows. **(A)** Distributions of predicted human-archaic hominin ancestral (orange) and expected human-archaic hominin ancestral (gray) 3D divergence for an example highly divergent window. Dotted lines represent the mean of their respective distributions. **(B)** Sharing of highly divergent windows among 1KG super-populations. Bars indicate the number of highly divergent windows present in each combination of populations indicated by the dot matrix. Population combinations with fewer than 10 windows are not plotted; see **Figure S5** for the full plot. **(C)** Example predicted maps for two African Yoruba individuals at the example window, one with low 3D divergence from the ancestor (NA18522; 3D divergence = 0.0002) and one with predicted high divergence (NA18486; 3D divergence = 0.031). The predicted maps are scaled to 10 kb resolution to be comparable to the resolution of the experimental Hi-C maps. **(D)** Experimentally determined Hi-C contact maps for this example window for the two Yoruba individuals. These experimental maps confirm the predicted high 3D divergence and contact pattern differences.

### 2.8 *In silico* mutagenesis reveals that multiple SNVs contribute to common 3D genome variation

To identify the variants underlying the differences observed in each 3D divergent window, we performed *in silico* mutagenesis. *In silico* mutagenesis is a computational technique that uses the ability of Akita to rapidly make predictions on any input sequence to identify and interpret potential causal variants. First, we extracted 616,222 very common (non-ancestral allele frequency > 10%) 1KG SNVs from the 392 divergent windows. We focused on common variants because large numbers of individuals have divergent 3D contact patterns in these windows. We inserted these variants one-by-one into the human-archaic hominin ancestral genome and used Akita to generate chromatin contact predictions for the mutated sequences in each window. Next, we calculated 3D divergence between the ancestral and mutated contact maps (**Figure 6A**) and quantified the effect of each SNV as the 3D divergence it produces from the ancestral map divided by the maximum 3D divergence between a modern human from ancestral for the window.

**Figure 6:**
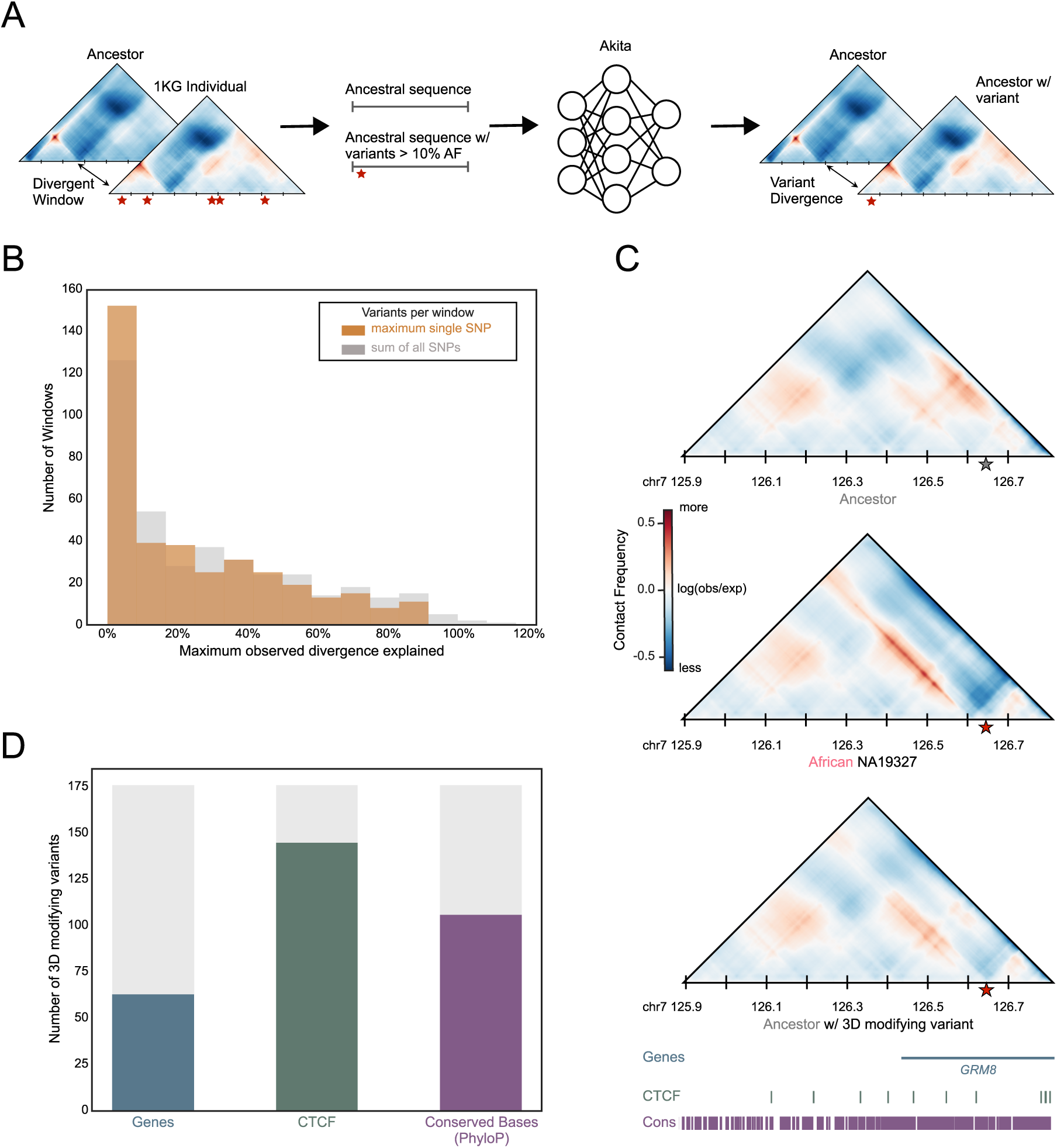
Most common divergent windows cannot be explained by a single nucleotide variant. **(A)** We used *in silico* mutagenesis to identify SNVs that likely contribute to 3D genome differences in highly divergent windows. First, we extracted common (non-ancestral AF > 10%) 1KG SNVs from the 392 windows with significantly greater 3D divergence across individuals than expected. We inserted the variants one-by-one into the human-archaic hominin ancestral genome and used Akita to generate chromatin contact predictions for the mutated sequences. Next, we calculated 3D divergence between the ancestral and mutated contact maps. Variants that produce greater than 20% of the maximum observed divergence in the window were designated 3D-modifying variants (N = 176). **(B)** Distribution of single SNV effects for the maximally disruptive SNV per window (gray) and for the linear sum of all SNV effects (orange). SNV effects are calculated as the percent of maximum divergence in a window between any 1KG individual and the ancestor that is observed in the mutated map. **(C)** Example SNV that recapitulates some, but not all, of the observed divergence from ancestral in a 3D divergent window. The tracks below the contact map show locations of genes (blue), CTCF binding sites (green) and phastCons elements (purple). **(D)** Number of the 176 3D modifying variants that are in CTCF binding peaks, genes, and conserved bases (phyloP).

A single SNV is not sufficient to explain the 3D divergence observed in most of these windows. For example, the maximum divergence explained by a SNV for each window is less than 10% of the overall 3D divergence(**Figure 6B**, orange) in more than 40% of windows. We designated the 176 variants that explain greater than 20% of the maximum observed divergence in a window as “3D-modifying variants”. We also find that summing the individual effects of all SNVs in a window does not recover substantially more of the observed 3D divergence from ancestral (**Figure 6B**, grey). This suggests that the divergence is not simply the result of additive combinations of the effects of common SNVs. To illustrate one of the strongest 3D-modifying variants, a SNV on chromosome 7 decreases the strength of an insulating region, causing overall structure in the window to be much less defined (**Figure 6C**). This SNV explains 38% of the 3D divergence between an African individual and the ancestor.

We quantified the number of 3D modifying variants overlapping CTCF peaks, genes, and conserved bases as called by phyloP (**Figure 6D**) (Pollard et al., 2010). 82% of 3D modifying variants are found in CTCF binding sites or and 60% are in conserved loci. Conversely, only 36% are found within genes. Our results suggest that the 3D-modifying variants identified exert a nuanced influence, as the maximum impact common SNV for each window contributes modestly to the predicted divergence. Furthermore, these variants predominantly occur at CTCF binding sites and conserved loci, rather than within genes. This underscores their potential significance in sculpting the 3D genomic architecture, especially considering the role of 3D chromatin contact in constraining sequence evolution.

### 2.9 26% of the genome has rare 3D genome variation

In the previous section, we investigated windows in which there was common variation in 3D chromatin contact patterns between individuals. We also observed a high occurrence of rare 3D genome variation—where one or a small number of individuals differ from a prevalent contact pattern. To discern underlying patterns in windows with rare 3D divergence, we implemented a classification scheme based on clustering contact maps. Strikingly, the most prevalent pattern was a single individual harboring rare variation that distinguished them from the remainder of the cohort (**Figure 7A**). This distinctive pattern was observed in approximately 26% of the windows (N = 1,251). Furthermore, the majority of windows exhibiting rare variation were primarily driven by an individual of African ancestry, characterized by substantial divergence from all other individuals in the study cohort (**Figure 7B**). The prevalence of single individuals exhibiting rare variation in a substantial proportion of windows underscores the potential of individual-specific genomic alterations to shape 3D genome architecture. Additionally, the prominent contribution of individuals of African descent to windows with rare variation highlights the importance of considering diverse genetic backgrounds when studying 3D genomic diversity.

**Figure 7:**
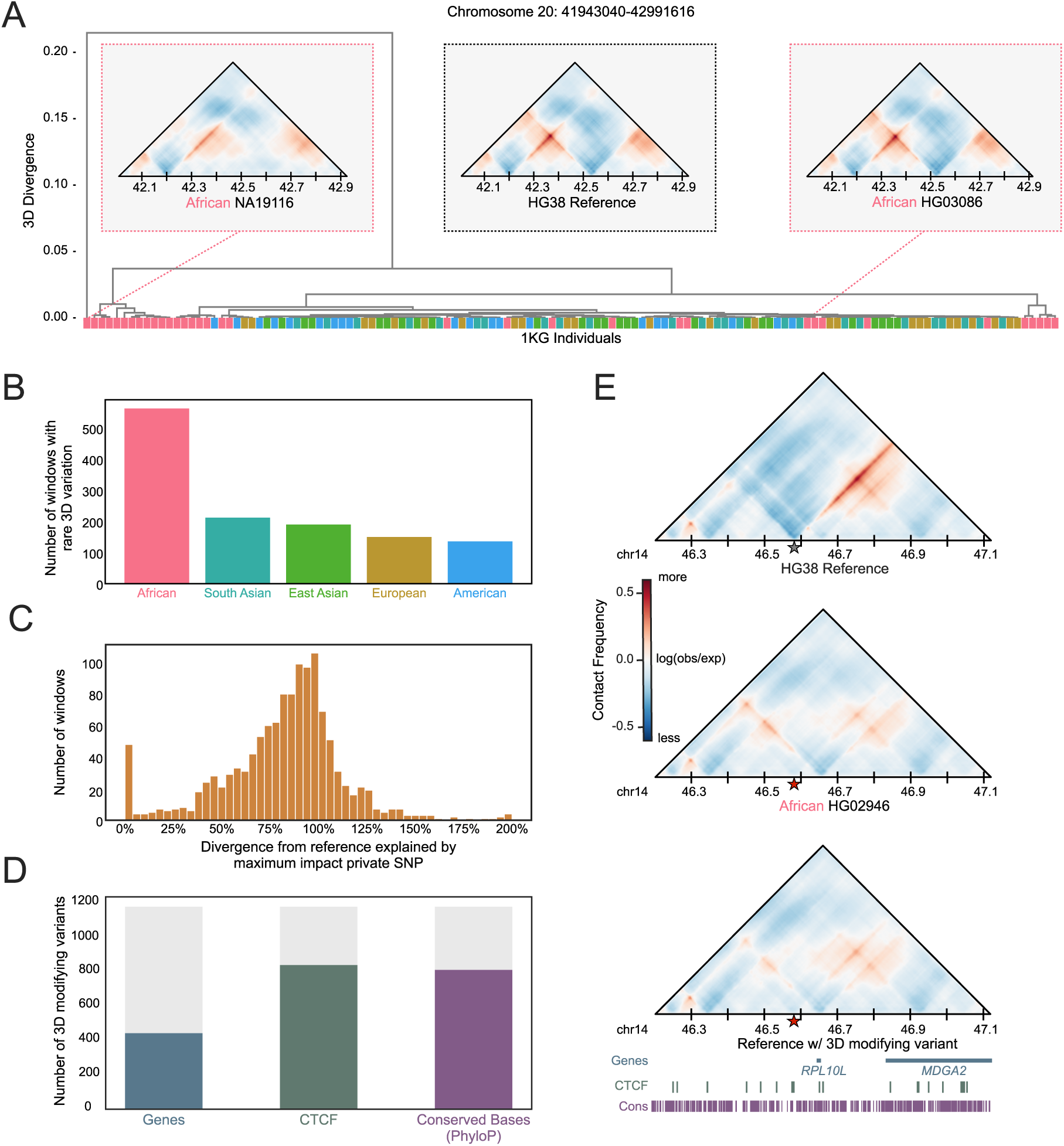
Genomic windows with rare 3D variation are common. **(A)** Tree of individuals based on 3D divergence for an example window where one individual is highly divergent from all others. The three contact maps show the patterns in the divergent individual, an individual from the same continental population, and the hg38 human reference. **(B)** Number of windows with rare 3D divergence stratified by continental origin of the rare individual. In total, 26% of windows in the genome have a rare divergent 3D contact pattern. **(C)** Distribution of single SNV effects for the maximally disruptive SNV per window. SNV effects are calculated as the percent of maximum divergence observed between a 1KG individual and the hg38 reference for a given window that is observed in the mutated map. **(D)** Number of the 3D modifying variants that are within CTCF binding peaks, genes, and conserved bases (phyloP). **(E)** Example of a single SNV that recovers the observed divergence in an individual with rare 3D variation when place into the reference sequence. The tracks below the contact map show the locations of genes (blue), CTCF binding sites (green) and phastCons elements (purple).

### 2.10 Rare 3D genome variation is usually the result of a single large-effect variant

To identify the variants contributing to the most prominent 3D differences in windows with rare 3D variation, we used *in silico* mutagenesis to test rare SNVs in the windows with rare 3D variation. We selected 59,797 variants that are private to the highly divergent individual (in the context of the 130 individuals used for the clustering analysis) to be inserted these one-by-one into the hg38 human reference genome and calculated 3D divergence between the reference and mutated contact maps (**Figure 6A**). We then quantified the effect of a SNV by calculating the percentage of the divergence between the highly divergent individual from the reference maps that is generated by inserting the SNV alone into the reference sequence.

In contrast to cases of common 3D divergence, the maximum explanatory SNV for each window often generates at least 100% of the observed divergence from reference (**Figure 7C**). In cases in which the divergence is greater for the single SNV, it suggests that other variants in the individual reduce the divergence. We identify 1,177 variants that explain at least 20% of the 3D divergence between the rare individual and the reference genome. 71% of these 3D modifying variants are found in CTCF binding sites or and 69% are in conserved loci. Conversely, only 38% are found within genes. To illustrate this pattern, we highlight an example SNV that decreases the strength of an insulating region, causing overall structure in the window to be much less defined (**Figure 7D**). This SNV explains 78% of the 3D divergence between an African individual and the ancestor. In contrast to our results in 3D divergent windows, these results suggest that rare 3D variation is often caused by a single, strongly 3D modifying variant.

## 3 Discussion

Our study explores the interplay between genetic sequence variation and 3D chromatin contact using machine learning to predict 3D chromatin contacts (Fudenberg et al., 2020) for thousands of diverse modern humans (1000 Genomes Project Consortium et al., 2015). Quantifying 3D chromatin contact on this scale is necessary to capture its variation across humans, and given the logistical and technical challenges of generating high-resolution Hi-C data at population-scale, currently this would not be possible without computational methods. The population-level perspective provided by our dataset enabled us to make several novel observations not seen in previous small-scale studies of human 3D chromatin contact diversity (Li et al., 2023).

### 3.1 3D chromatin contact divergence vs. sequence divergence

Our results show that 3D chromatin contact divergence follows similar genome-wide trends as sequence divergence. For example, African populations exhibited consistently higher average 3D divergence in comparison to other populations, which corresponds to their greater sequence diversity (1000 Genomes Project Consortium et al., 2015). However, the correlation between 3D chromatin contact similarity and sequence divergence (*R*^2^ = 0.31) is only moderate, suggesting the existence of differing influences and regulatory mechanisms shaping the interplay between sequence divergence and 3D genome organization across diverse individuals. Indeed, quantification of local window-specific divergence showed that 3D contact map variation in most genomic regions is shared across populations, and no windows have contact map patterns that stratify by population. Moreover, it revealed that rare 3D contact variation is common—26% of windows have an individual with a rare divergent contact pattern.

### 3.2 Influence of 3D chromatin contact on sequence evolution and functional constraint

We also found that the observed 3D divergence between modern humans and the human-archaic hominin ancestor is significantly less than anticipated based on observed sequence divergence. This suggests that constraint imposed by the pressure to maintain 3D chromatin contacts shaped sequence divergence during recent human evolution. The findings are consistent with prior studies indicating a depletion of variation at key 3D genome determining elements (Fudenberg and Pollard, 2019; Krefting et al., 2018; McArthur and Capra, 2021; McArthur et al., 2022; Sauerwald et al., 2020; Whalen and Pollard, 2019) and suggest that the preservation of 3D chromatin contact contributes to sequence constraint in human evolution. By comparing the observed and expected 3D divergence derived from sequence divergence, we underscore the potential role of 3D genome organization in influencing recent human sequence evolution. Examining local patterns of 3D divergence along the genome revealed substantial variability, indicating varied tolerance for 3D genome divergence. We found that regions exhibiting elevated 3D divergence consistently had reduced gene content, fewer CTCF binding sites, and fewer conserved bases than other genomic windows. These results are consistent with previous work that investigated two cell lines and found variation along chromosomes that correlates with compartment, GC content, transcription rate and repeat element prevalence (Gunsalus et al., 2023a). This pattern underscores the importance of maintaining 3D chromatin contacts, especially in regions with many functional elements, on shaping evolutionary pressures.

### 3.3 *In silico* mutagenesis identifies SNVs likely to drive 3D divergence

Another strength of the sequence-based machine learning approach is that it enables rapid screening of the effects of individual genetic variants in different genetic backgrounds (Brand et al., 2023; Gunsalus et al., 2023b; McArthur et al., 2022). We used this *in silico* mutagenesis to unravel the influence of SNVs on 3D genome variation. We discovered that the 3D divergence in windows with common 3D variation was rarely the result of the independent additive effects of common SNVs. Instead, our analyses suggest that combinations of SNVs likely interact to produce much of the common variation in the 3D genome. In contrast, for windows with only rare 3D variation, a single, high-impact variant was often sufficient to produce the observed divergence. This suggests that individual variants with strong impacts on 3D contact are rarely tolerated at high frequencies. However, the 3D-modifying variants observed in both types of windows predominantly influenced crucial functional sites such as CTCF binding sites and evolutionarily conserved loci. The sharp contrast in the nature of variant contributions to common and rare 3D variation underscores the intricate mechanisms governing 3D chromatin contact and its variation.

### 3.4 Machine learning addresses challenges with experimental Hi-C data

Traditional Hi-C experiments often compromise resolution for coverage, resulting in representations that lack finer details pivotal for understanding 3D genome architecture at the scale of differences observed between healthy individuals. This drawback limits our ability to capture subtle but potentially functional chromatin interactions, impeding comprehensive genomic analysis. To overcome these limitations, our study harnesses an accurate machine learning prediction model called Akita. Akita demonstrates robust performance in generating local 3D contact patterns from DNA sequences at a higher resolution (2 kb), enabling a finer depiction of chromatin interactions that compensates for experimental limitations (Fudenberg et al., 2020). Our findings showcase Akita’s efficacy in predicting 3D chromatin architecture not only in European-derived cell lines, its original training data, but also in diverse populations, particularly among African individuals. This ability to perform consistently across diverse populations is critical, as it allows us to investigate chromatin organization in populations where Hi-C data is limited. Our research thereby offers a more comprehensive view of the 3D genome landscape, crucial for understanding chromatin organization and its functional implications. Our computational approach addresses the limitations of low-resolution Hi-C and enables the exploration of 3D chromatin contact in a broader range of individuals with available genome sequences. Leveraging Akita’s versatility, we extend the analysis to encompass thousands of individuals from diverse populations, transcending the boundaries of experimental resolution.

### 3.5 Limitations

While our study increases our understanding of chromatin contact variation, it is important to acknowledge several limitations. First, the current constraints in Hi-C data quality and quantity limit the resolution of our analyses and experimental validation. The machine learning models that enable this work are limited in performance by the quality of the available training data. Second, while we validate example predictions with experimental Hi-C data, our results are based on machine learning models that do not have perfect performance. We are confident in our conclusions, but moving forward, we envision close integration between computational predictions and new experimental data for validation and discovery of the intricacies of 3D chromatin contact. Additionally, the 1KG dataset, while extensive, does not encompass the entirety of human genetic diversity. Specifically, the African individuals included in 1KG do not capture more deeply divergent African lineages; expanding to additional datasets would increase the scope of genetic diversity covered (Fan et al., 2023; Mallick et al., 2016). Hence, future studies should aim to incorporate a wider array of populations to provide a more comprehensive understanding of the interplay between 3D chromatin contact and genetic sequence divergence. Our study is also solely focused on SNVs, excluding structural variants, which have been shown to contribute to 3D chromatin contact differences (Norton and Phillips-Cremins, 2017; Sánchez-Gaya et al., 2020; Spielmann et al., 2018). Additionally, our analysis did not explore the potential impact of differences between various cell-types, which could influence the observed 3D chromatin contact patterns. Further validation of the predicted relationship between sequence variation, 3D chromatin contact, and functional implications presented in this study necessitates increased data resolution, depth and cell-type coverage. Future efforts to expand Hi-C data resolution and availability are essential to comprehensively understand the mechanisms and variation of chromatin organization and its functional outcomes.

### 3.6 Conclusions

Our study uses machine learning to map the relationship between genetic sequence variation and 3D chromatin contact across diverse human populations. Our findings pave the way for future research exploring the mechanisms governing chromatin organization and its functional implications in disease and evolution.

## 4 Methods

### 4.1 Modern human and ancestral genomes

All genomic analysis was conducted using the GRCh38 (hg38) genome assembly and coordinates. Genomic variation within modern humans came from 1000 Genomes Project (1KGP), Phase 3 from (1000 Genomes Project Consortium et al., 2015). The ancestral human genome was extracted using ancestral allele calls for each position in the tree sequence from (Wohns et al., 2022). Tree sequences are an efficient data format for representing the ancestral relationships between sets of DNA sequences and were analyzed using tskit (Kelleher et al., 2018). We constructed full-length genomes for each individual based upon the genotyping information in their respective VCF file. We treated all individuals as if they were homozygous (pseudo-haploid). We built each individual genome using GATK’s Fasta Alternate Reference Maker tool (Van der Auwera and O’Connor, 2020). If an individual had an alternate allele (homozygous or heterozygous), we inserted it into the reference genome to create a pseudo-haploid, or “flattened” genome for each individual. To maintain the necessary consistent window and overlap size required by Akita, we included SNVs but not SVs in these genomes.

### 4.2 3D chromatin contact prediction with Akita

After the genomes were prepared, we input them into Akita for predictions using a *∼*1 Mb sliding window (1,048,576 bp) overlapping by half (e.g. 524,288-1,572,864, 1,048,576-2,097,152, 1,572,864-2,621,440). Although Akita was trained simultaneously on Hi-C and Micro-C across five cell types in a multi-task framework to achieve greater accuracy, we focused on predictions in the highest resolution maps, human foreskin fibroblast (HFF) as in McArthur et al., 2022. Akita considers the full window to generate predictions, but the resulting predictions are generated for only the middle 917,504 bp. Each contact map is a prediction for a single individual, and each cell represents physical 3D contacts at 2,048 bp resolution. The value in each cell is log2(obs/exp)-scaled to account for the distance-dependent nature of chromatin contacts. For all analyses, we only considered windows with 100% coverage in the hg38 reference genome for a total of 4,873 autosomal windows. Fudenberg et al., 2020 provides further details on the CNN architecture and training data used.

### 4.3 3D and sequence genome comparisons

After predictions were made on all 1 Mb windows for all individuals, we compared the resulting predictions using mean-squared error and Spearman and Pearson correlations. All measures are scaled to indicate divergence: higher indicates more difference while lower indicates more similarity. In the main text we transformed the Spearman’s rank correlation coefficient (1-*ρ*) to describe 3D divergence (Gunsalus et al., 2023b). Some analyses compared 3D genome divergence with sequence divergence. To calculate the sequence divergence between two individuals, we counted the proportion of bases at which the two individuals differ in the 1 Mb window. This was done only for windows with 100% coverage in hg38, as with the 3D chromatin contact predictions.

### 4.4 Empirical distribution of expected divergence

We generated genomes with shuffled nucleotide differences to compute the expected 3D divergence in a window given the observed sequence divergence. This approach was adapted from McArthur et al., 2022. We matched these shuffled differences to the same number and trinucleotide context of the observed sequence differences between an individual genomes from each population (HG03105 [African], HG01119 [American], NA06985 [European], HG00759 [East Asian], HG03007 [South Asian]) and the inferred ancestral genome. For each 1 Mb window of the genome (N = 4,873) we generated 500 shuffled sequences. We calculated an empirical distribution of expected 3D divergence from comparing the contact maps of the shuffled sequences with the ancestral sequence. Finally, we compared the average expected 3D divergence from this distribution to the observed ancestral-modern 3D divergence.

### 4.5 3D genome divergence vs. functional/conserved elements

Conservation and functional genome annotations were obtained from publicly available data sources. Gene annotations are from GENCODE version 24 (Frankish et al., 2019). CTCF binding sites were determined through ChIP-seq analyses from ENCODE (“An Integrated Encyclopedia of DNA Elements in the Human Genome” 2012; Davis et al., 2018). We downloaded all CTCF ChIP-seq data with the following criteria: experiment, released, ChIP-seq, human (hg38), all tissues, adult, BED NarrowPeak file format. We excluded any experiments with biosample treatments. Across all files, CTCF peaks were concatenated, sorted, and merged into a single file, merging overlapping peaks into a single larger peak. We quantified the number of CTCF ChIP-seq peaks per genomic window (peaks per window) and the number of CTCF peak base pairs overlapping each window (base pairs per window), Evolutionary constraint was quantified by PhastCons as described above. The PhastCons elements (Siepel et al., 2005) were intersected with 1Mb genomic windows, partitioned by 3D divergence. The overlap quantification is the number of PhastCons base pairs per boundary regardless of score (base pairs per window). Convserved base pairs were identified by PhyloP (Pollard et al., 2010), using PhyloP scores downloaded from the UCSC Genome Table Browser (https://genome.ucsc.edu/cgibin/hgTables)

We used the pybedtools wrapper for BEDtools (Dale et al., 2011; Quinlan and Hall, 2010) to perform intersections of genomic regions for the above annotations (genes, CTCF peaks, PhastCons) with the 1Mb windows used for Akita predictions. These windows were stratified by mean 3D divergence from ancestral for all 1KG individuals and by the difference in the mean of the observed distribution of 3D divergence from the expected as described above.

### 4.6 Shared divergent windows across populations

The top 10% of windows for each population were chosen based on the mean 3D divergence from the ancestral for all individuals in the respective populations. Overlap was calculated using a python implementation of UpSet plots, a tool to visualize set overlaps (Lex et al., 2014; Nothman, 2023).

### 4.7 Hierarchical clustering of 3D chromatin contact maps

A subset of 130 1KG individuals, chosen at random to represent 5 individuals from each population, were compared in a pairwise fashion across all 4873 genomic windows. Pairwise 3D divergence score matrices for each 1Mb window were used to perform hierarchical clustering on these individuals, plus the human-archaic hominin and human-chimp ancestral genomes, using the hierarchical clustering functionality from scipy with complete linkage (Virtanen et al., 2020). The clustering generated dendrogram “trees” that describe the relationships between individuals. The Python API for ETE ToolKit (Huerta-Cepas et al., 2016) was used to identify any trees that are monophyletic for a given population, meaning that any population clustered entirely and exclusively together. To establish support for known population patterns based on the 3D divergence trees, we generated a baseline tree representing the sequence similarity of two inferred ancestors and 1KG individuals from all populations but the Americas super population and African American Southwest (ASW) population, which exhibit substantially more admixture than other populations (1000 Genomes Project Consortium et al., 2015; Bergström et al., 2020; Duda and Jan Zrzavý, 2016; Gravel et al., 2011; Li et al., 2008). To calculate support for the branches of this tree ASTRAL (Rabiee et al., 2019; Zhang et al., 2018) was used, treating the window based tree as ‘gene trees’ and the baseline tree as a ‘species’ tree.

### 4.8 *In silico* mutagenesis

We identified individual variants contributing to 3D divergence among commonly 3D divergent windows using an *in silico* approach (**Figure 6A**). We identified common non-ancestral alleles (AF>10%) among the 1KG individuals, consisting of 616,222 unique variants in 392 genomic windows. For each variant-window pair, we inserted the variant into the ancestral sequence for that window and calculated the 3D divergence between the ancestral map and the ancestral with variant map. “3D-modifying variants” were defined as variants (add criteria here).

We calculated the effects of 3D-modifying variants by calculating “explained divergence” by dividing the 3D divergence for the variant by the maximum ancestral to 1KG comparison for the window. Values near zero indicate that the 3D-modifying variant explains minimal divergence among the observed comparisons, while values near one indicate the variant explains most of the divergence among observed comparisons. Values greater than one indicate that variant creates more 3D divergence than observed among any ancestral to 1KG comparison, suggesting that other variants may “buffer” against the variant’s effect.

We also applied our *in silico* mutagenesis approach to rare variants private to the highly divergent individual in each of 1,251 windows with rare 3D variation. Private variants were defined at positions where only the focal individual carried a copy of the alternate allele. This was done only in the context of the 130 individuals used for clustering analysis. The individual of interest had a genotype with at least one non-reference allele, whereas all others were fixed for the reference allele. We considered 59,797 variants in the 1,251 windows. In this case explained divergence was calculated with respect to the hg38 reference genome as this analysis focuses on within human variation.

### 4.9 Analysis of experimental HiC data

Preprocessed cooler files were downloaded from the 4DN Data Portal (https://data.4dnucleome.org) and analyzed at 10 kb resolution. Visualization was done using custom code adapted from Brand et al., 2023, Gunsalus et al., 2023b and Fudenberg et al., 2020.

### 4.10 Significance reporting

The machine used to run analyses had a minimum value for representing floating numbers of 2.2250738585072014 *×* 10^−308^. Therefore, we abbreviate values less than this as 2.23 *×* 10^−308^.

### 4.11 Data availability

The publicly available data used for analysis are available in the following repositories: 1KG VCFs are available at (ftp.1000genomes.ebi.ac.uk/vol1/ftp/data_collections/1000_genomes_project/release/20190312_biallelic_SNV_and_INDEL/)(1000 Genomes Project Consortium et al., 2015).

CTCF-bound open chromatin candidate cis-regulatory elements (cCREs) in all cell types (https://screen.encodeproject.org/ *>* Downloads *>* Download Human CTCF-bound cCREs). phast-Cons elements and PhyloP scores were retrieved from the UCSC Genome Browser (https://hgdownload.soe.ucsc.edu/goldenPath/hg38/database/phastConsElements100way.txt.gz, https://hgdownload.soe.ucsc.edu/goldenPath/hg38/database/phyloP100way.txt.gz).

Experimental HiC available at the 4D nucleome data portal (https://data.4dnucleome.org/browse/?dataset_label=Hi-C+on+lympoblastoid+cell+lines+from+1000G+individuals&experiments_in_set.experiment_type.experiment_category=Sequencing&experimentset_type=replicate&type=ExperimentSetReplicate)

### 4.12 Code availability

All code used to conduct analyses and generate figures is publicly available on GitHub (https://github.com/egilbertson-ucsf/3DGenome-diversity). Akita is available from the basenji repository on GitHub (https://github.com/calico/basenji/tree/master/manuscripts/akita).

## 4.13 Acknowledgements

This work was supported by the National Institutes of Health (NIH) General Medical Sciences Institute award R35GM127087 to JAC, NIH National Heart, Lung, and Blood Institute award U01HL157989 to KSP, and NIH National Human Genome Research Institute award F30HG011200 to EM. The work as also supported by funds from the Gladstone Institutes and the Bakar Computational Health Sciences Institute.

This work was conducted in part using the resources of the Wynton High Performance Computer at the University of California San Francisco.

We thank Jian Ma’s lab for sharing SPIN State annotations for HFF cells.

We also thank members of the Capra and Pollard Labs who gave helpful feedback throughout this project.

## 4.14 Author Contributions

Conceptualization: ENG, CMB, EM, DCR, KSP and JAC Formal Analysis: ENG Visualization: ENG, CMB, and JAC Resources and Software: CMB, EM, DCR, SK Writing – Original Draft: ENG and JAC Writing – Review & Editing: ENG, CMB, EM, DCR, SK, KSP and JAC.

## 4.15 Competing interests

The authors declare no competing interests.

## Supplementary Information

### Supplemental Figures

**Figure S1:**
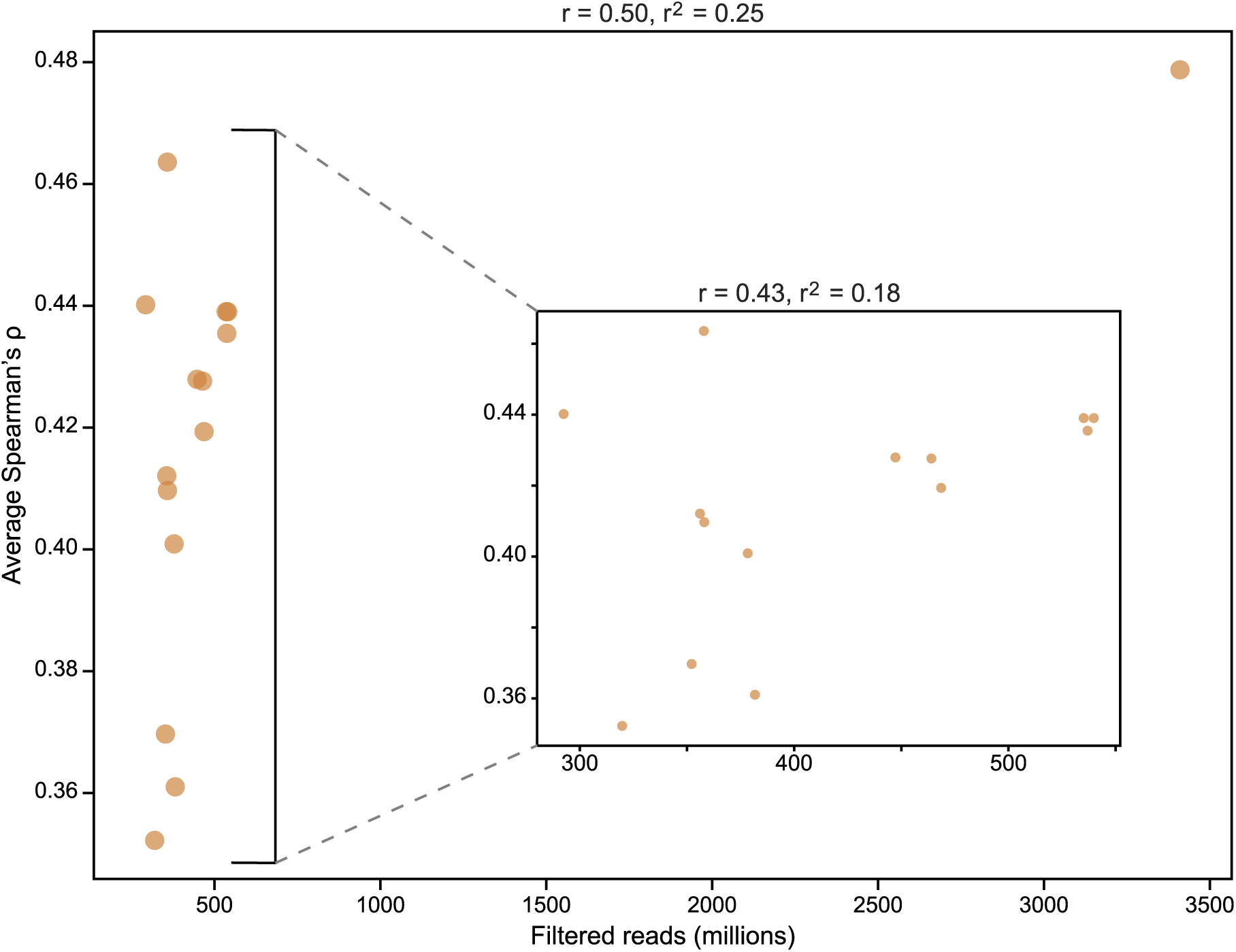
Correlation between read count and prediction accuary. Filtered read count in millions correlated with genome-wide average Spearman’s rho (predicted vs experimental) for 15 individuals for which we have experimental HiC data and Akita predictions at 10 kb resolution.

**Figure S2:**
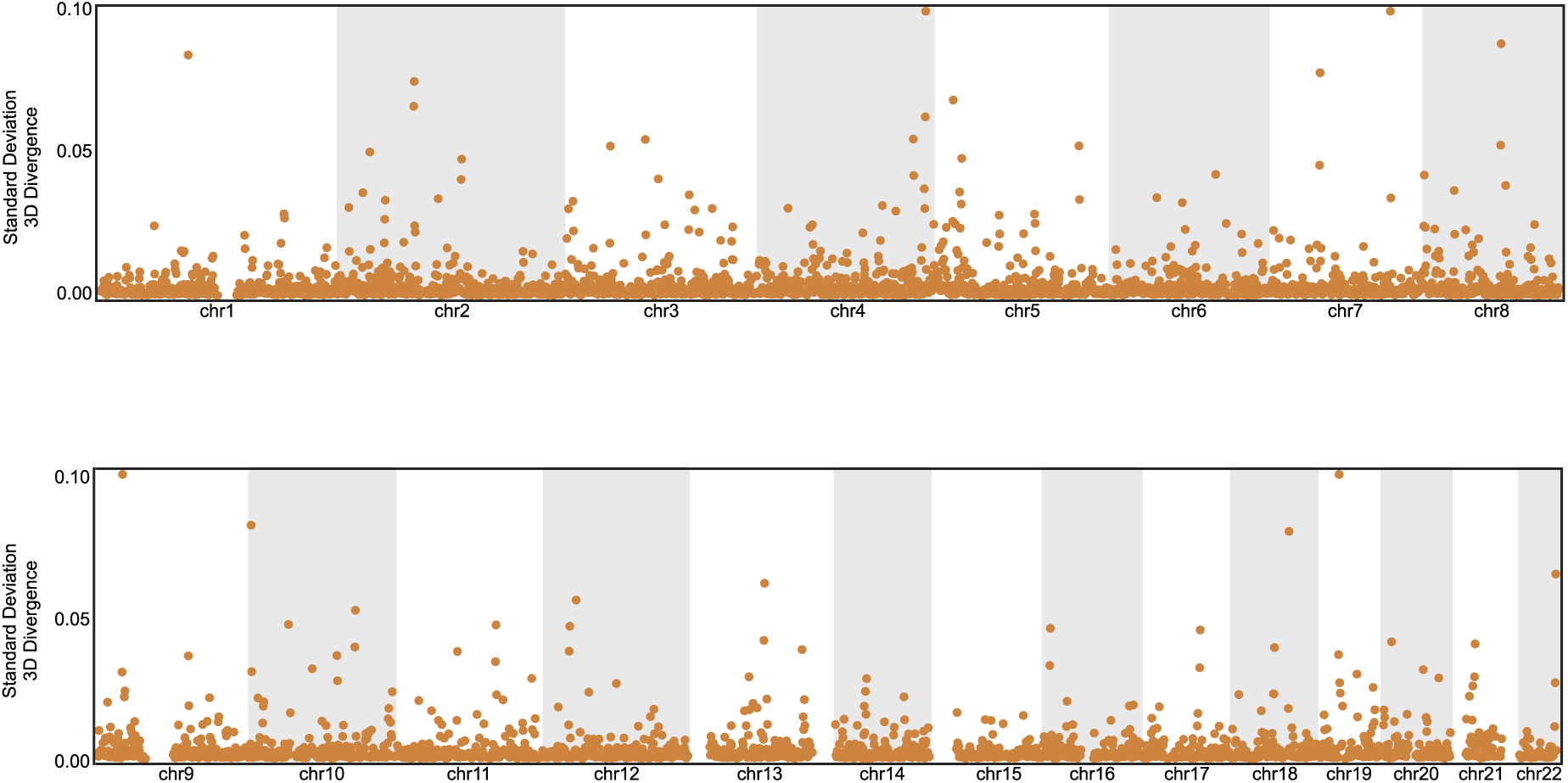
Genome-wide standard deviation of 3D divergence between all 1KG individuals. Standard deviation of genome-wide 3D divergence from the human-archaic hominin ancestor across 4,873 genomic windows of 2,448 individuals from 1KG. Each point represents the standard deviation of divergence of all individuals from the ancestral genome for a single genomic window. All points greater than 0.10 are clipped to 0.10 for visualization.

**Figure S3:**
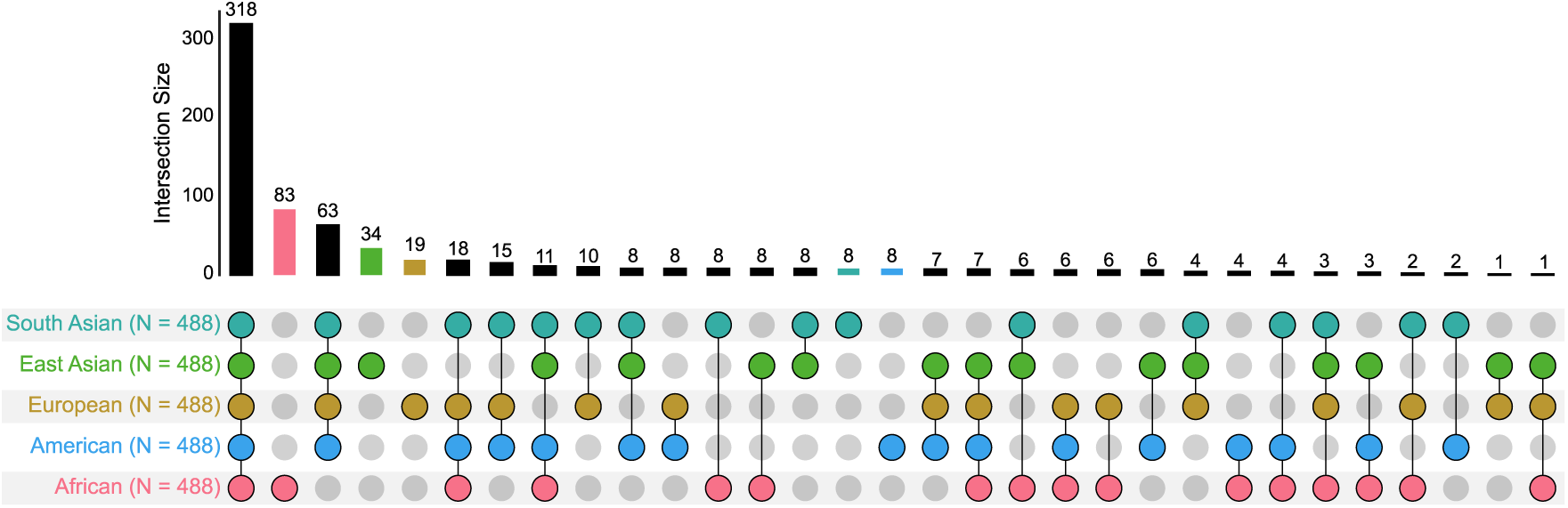
Upset plot of population of divergence for top 10% divergent windows. Unique and shared top 10% divergent windows among 1KG super-populations. Bars indicate the number of windows and the dot matrix indicates the populations represented by each set

**Figure S4:**
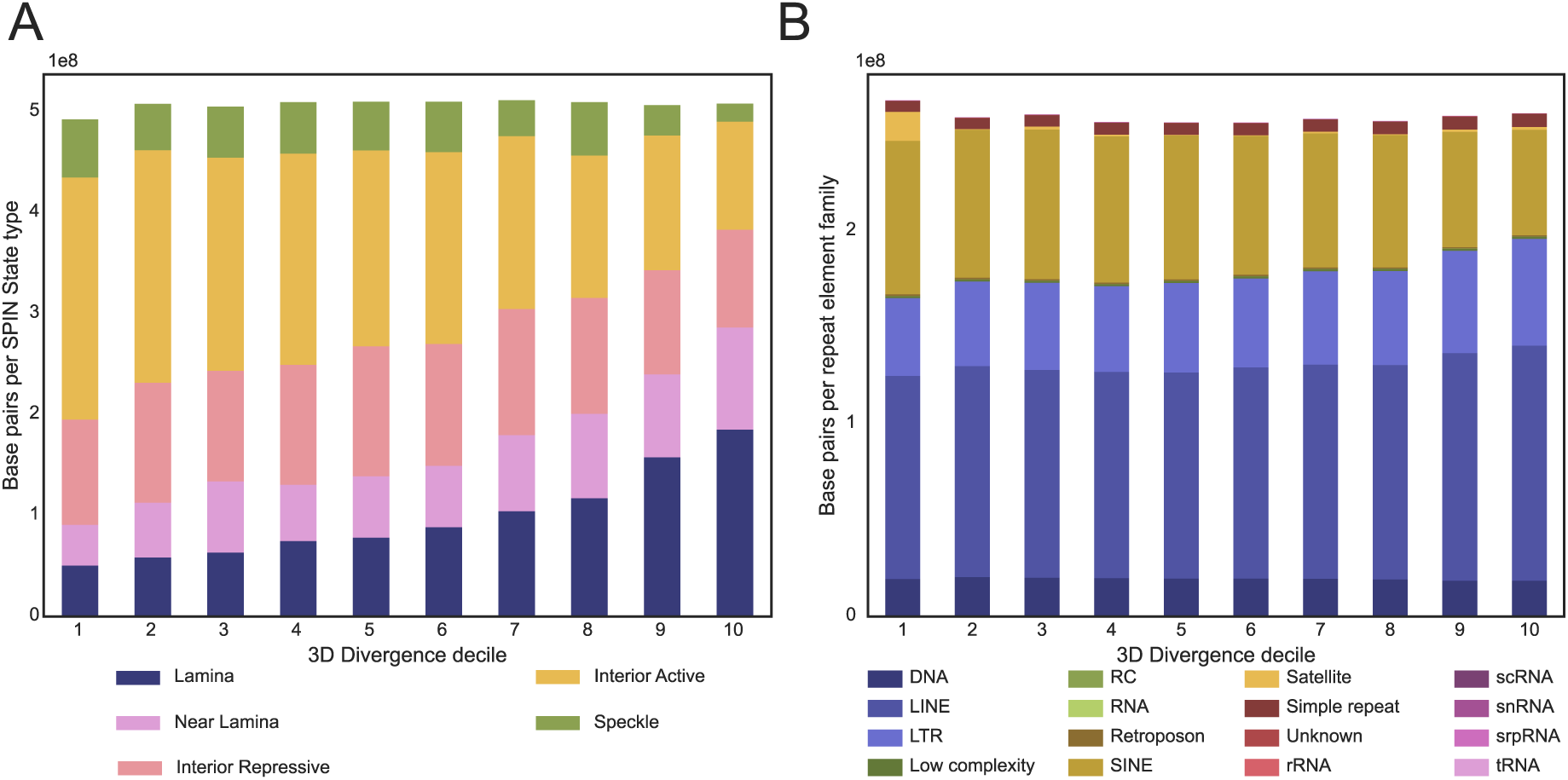
SPIN state and repeat element content across 3D divergence deciles. **(A)** SPIN states as in Wang et al., 2021, kindly provided for the HFF cell line by Jian Ma’s lab. The number of base pairs assigned to each SPIN state is shown according to presence each decile of 3D divergence. **(B)** The number of RMSK repeat element base pairs of each repeat family is shown according to presence each decile of 3D divergence.

**Figure S5:**
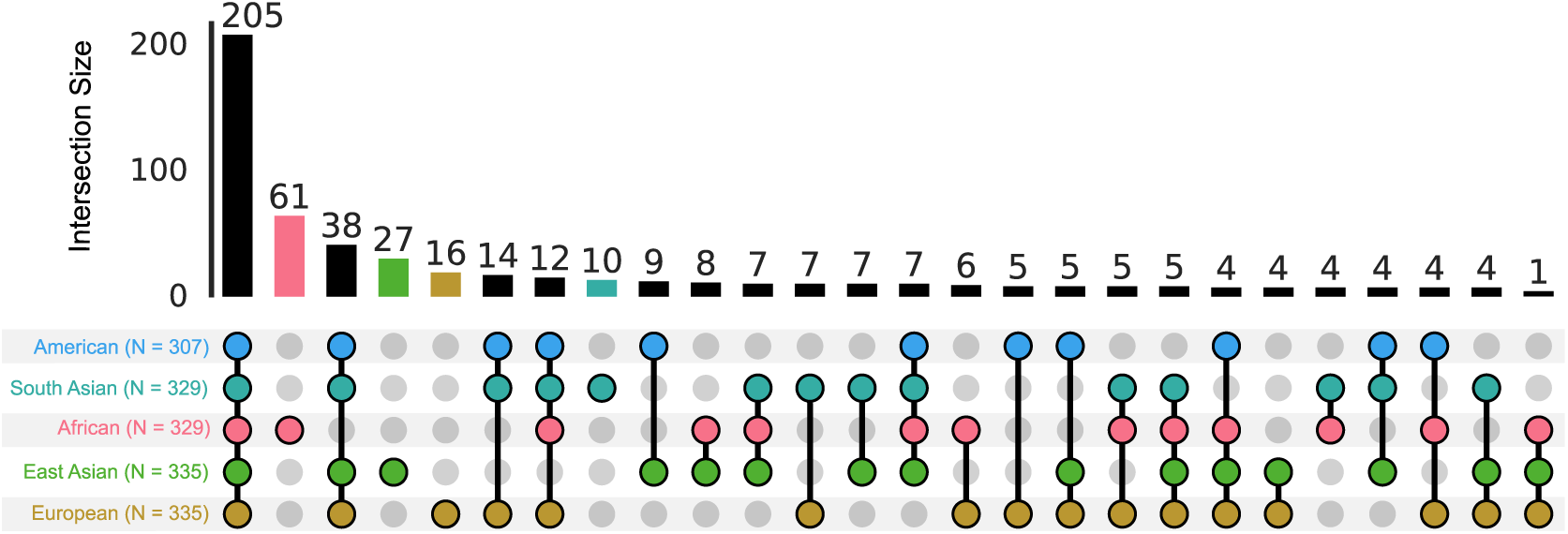
Full upset plot for more divergent than expected windows. Unique and shared divergent windows among 1KG super-populations. Bars indicate the number of windows and the dot matrix indicates the populations represented by each set.

## Notes

### Competing Interest Statement

The authors have declared no competing interest.

