## Supplemental Figures for "Machine learning reveals the diversity of human 3D chromatin contact patterns"

### Supplementary Information

#### Supplemental Figures

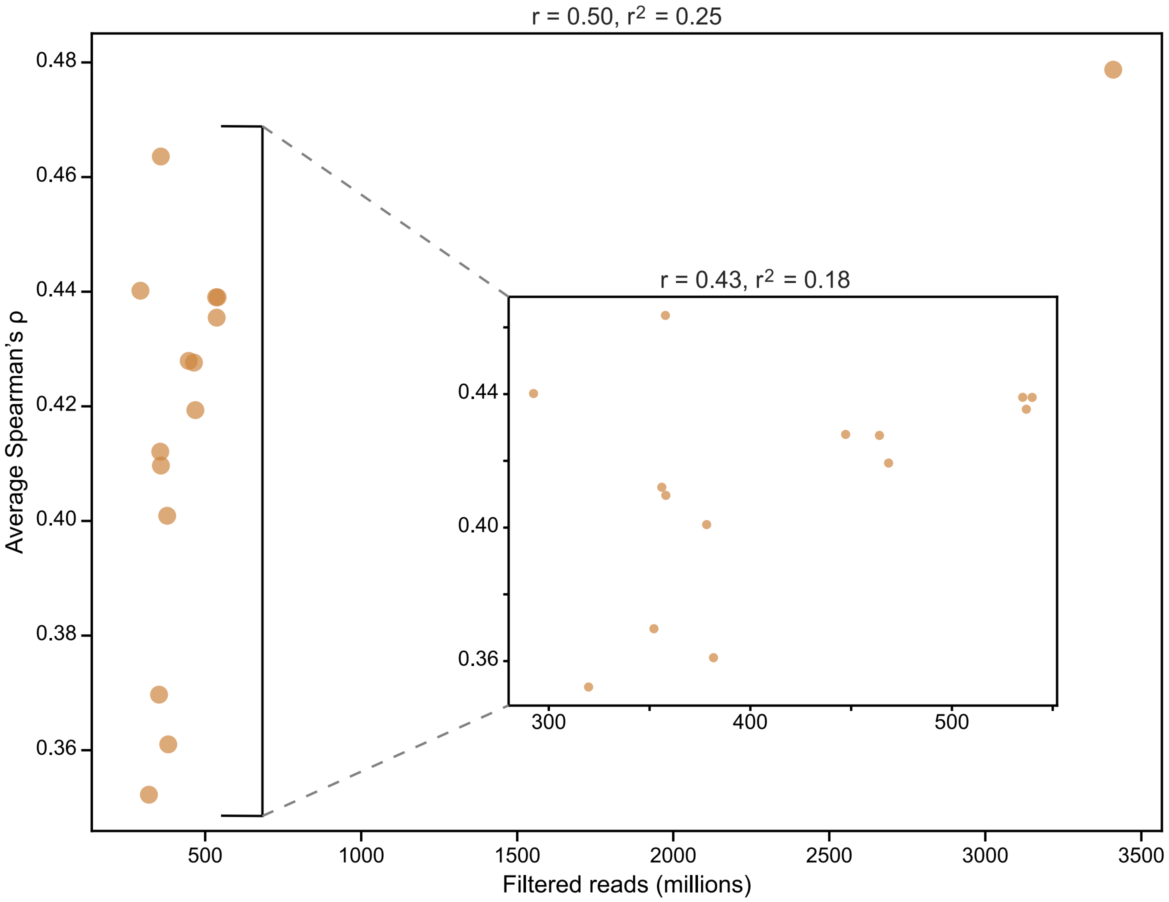


**Figure S1: Correlation between read count and prediction accuary.** Filtered read count in millions correlated with genome-wide average Spearman's rho (predicted vs experimental) for 15 individuals for which we have experimental HiC data and Akita predictions at 10 kb resolution.

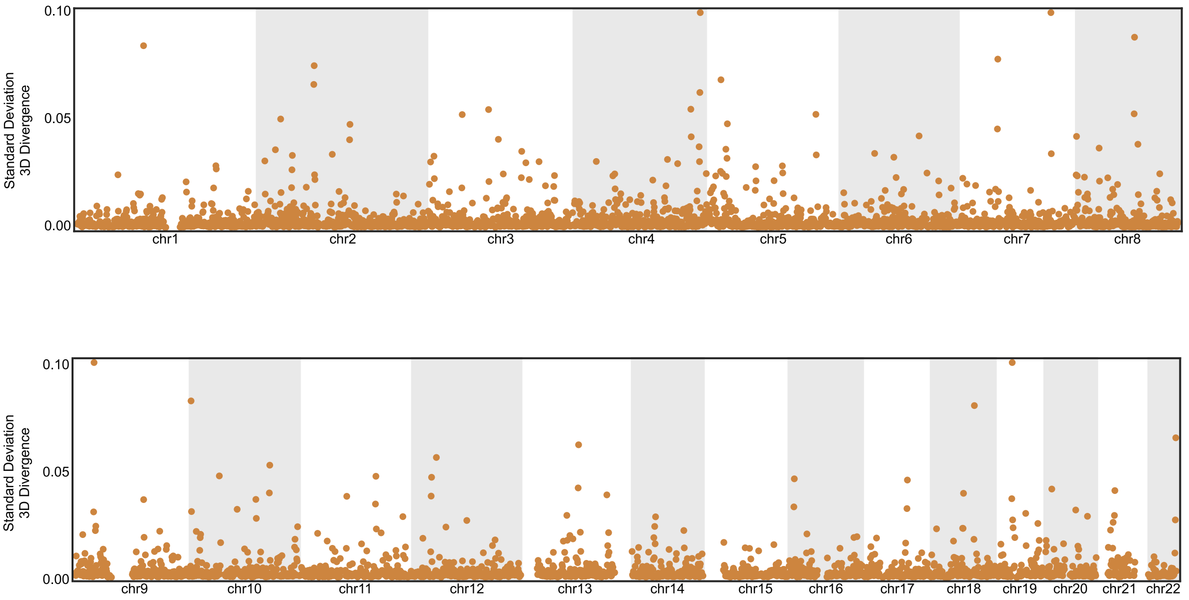


**Figure S2: Genome-wide standard deviation of 3D divergence between all 1KG individuals.** Standard deviation of genome-wide 3D divergence from the human-archaic hominin ancestor across 4,873 genomic windows of 2,448 individuals from 1KG. Each point represents the standard deviation of divergence of all individuals from the ancestral genome for a single genomic window. All points greater than 0.10 are clipped to 0.10 for visualization.

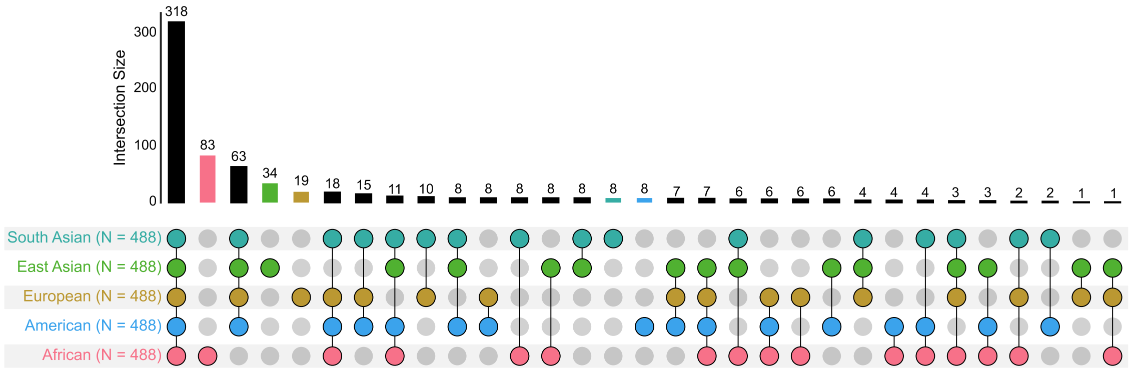


**Figure S3: Upset plot of population of divergence for top 10% divergent windows.** Unique and shared top 10% divergent windows among 1KG super-populations. Bars indicate the number of windows and the dot matrix indicates the populations represented by each set

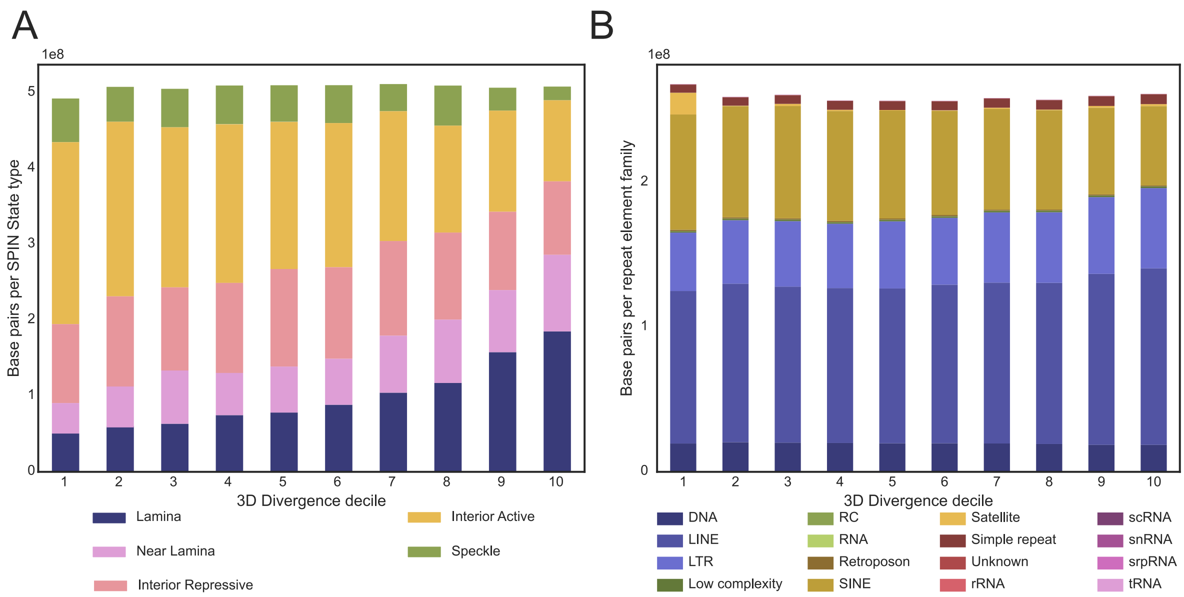


**Figure S4: SPIN state and repeat element content across 3D divergence deciles.** **(A)** SPIN states as in Wang et al., 2021, kindly provided for the HFF cell line by Jian Ma's lab. The number of base pairs assigned to each SPIN state is shown according to presence each decile of 3D divergence. **(B)** The number of RMSK repeat element base pairs of each repeat family is shown according to presence each decile of 3D divergence.

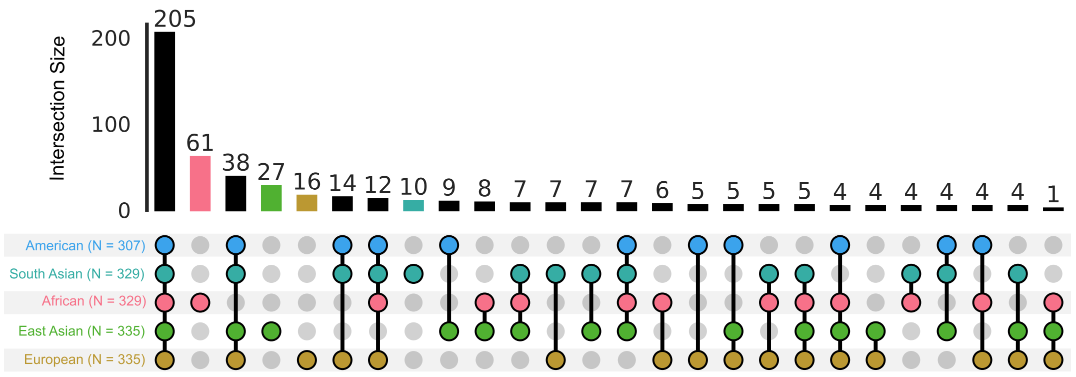


**Figure S5: Full upset plot for more divergent than expected windows.** Unique and shared divergent windows among 1KG super-populations. Bars indicate the number of windows and the dot matrix indicates the populations represented by each set.
